# Methylglyoxal modification of TrkB promotes synaptic plasticity and enhances resilience to chronic stress

**DOI:** 10.1101/435867

**Authors:** Ziyin Wu, Yingxue Fu, Yinfeng Yang, Chao Huang, Chunli Zheng, Zihu Guo, Zihao Yang, Xuetong Chen, Jinglin Zhu, Jinghui Wang, Xiaogang Li, Liyang Chen, Weiwei Zhou, Yangyang Chen, Jiangmei Wang, Yang Yang, Meng Jiang, Sushing Chen, Aiping Lu, Jianlin Liu, Yan Li, Shiguo Sun, Zhenzhong Wang, Wei Xiao, Yonghua Wang

## Abstract

The endogenous metabolite methylglyoxal (MGO) has recently emerged as a potential mediator of psychiatric disorders, such as anxiety and depression, but its precise mechanism of action remains poorly understood. Here, we find that MGO concentrations are decreased in the prefrontal cortex and hippocampus in rats subjected to chronic stress, and low-dose MGO treatment remarkedly enhances resilience to stress and alleviates depression-like symptoms. This effect is achieved by MGO’s promotion on the synaptic plasticity in prefrontal cortex and hippocampus. Both *in vitro* and *in vivo* experiments show that MGO provokes the dimerization and autophosphorylation of TrkB and the subsequent activation of downstream Akt/CREB signaling, which leads to a rapid and sustained expression of brain-derived neurotrophic factor (BDNF). We further demonstrate that MGO directly binds to the extracellular domain of TrkB, but not its intracellular domain. In addition, we also identify a natural product luteolin and its derivative lutD as potent inhibitors of Glyoxalase 1 and validate their antidepressant effects in chronic stress rat models. The antidepressant role of endogenous MGO provides a new basis for the understanding and therapeutic intervention design for stress-associated mental disorders.

## Introduction

Major depression and anxiety disorders are highly prevalent mental illness characterized by profound disturbances in emotional regulation of affected individuals ^1^. Chronic stress is a trigger for depression, and defects in the normal adaptivity to chronic stress can increase the risk of developing depression and anxiety disorder ^2^. The ability to cope with stressful situations, resilience, depending on the development of adequate behavioral and psychological adaptations to chronic stress, plays an essential role in modulating the development of and recovery from depression ^3, 4^. In view of the association between stress resilience and depression, identification of the neural substrates underlying stress resilience is critical for the development of therapeutic strategies for stress-related psychiatric diseases ^5, 6^.

Methylglyoxal (MGO) is an endogenous metabolite mainly generated in the glycolysis process ^7^. It is a highly reactive dicarbonyl aldehyde, which can react with protein arginine and/or lysine residues to form advanced glycation end-products (AGEs), and cause non-enzymatic, post-translational modifications (PTMs) of proteins ^8, 9^. High concentrations of MGO exert cytotoxic effects via evoking the production of reactive oxygen species ^10^, and have been implicated in pathologies of several diseases including diabetes, aging and neurodegenerative diseases ^11-13^. Nevertheless, under normal conditions, MGO can be efficiently detoxified by the glyoxalase system, mainly through Glyoxalase 1 (GLO1), an enzyme catalyzing the conversion of acyclic α-oxoaldehydes to corresponding α-hydroxyacids ^14^. Recently, MGO and GLO1 has been identified associated with anxiety- and depression-like behaviors in mice ^15, 16^. This suggests that MGO may play a potential role in the pathophysiology of depression. However, the cellular function of MGO at physiological concentration and its precise mechanism of action in depression still remain poorly understood.

In this study, we hypothesize that MGO could promote resilience to chronic stress. To test this, we constructed different stress models, and found that MGO levels are remarkedly decreased in the prefrontal cortex (PFC), hippocampus (HC) and also plasma in rats subjected to chronic stress, whereas low-dose MGO treatment remarkedly enhances resilience to stress and alleviates depression-like symptoms. This effect is achieved by MGO’s direct binding to the extracellular domain (ECD) of TrkB to stimulate its dimerization and autophosphorylation. This enables MGO to switch on the TrkB signaling pathways, which leads to a rapid and sustained expression of BDNF. MGO also effectively promotes the hippocampal neurogenesis in CMS rats. In addition, the modulation on MGO concentration by knockdown of GLO1 also exhibits antidepressant-like effects. The antidepressant role of endogenous MGO thus provides a new basis for the design of therapeutic interventions for major depressive disorder.

## Results

### MGO increases resilience to chronic stress and restores neurotransmitter levels

To investigate whether chronic stress leads to changes in MGO concentrations in the brain, we constructed a chronic mild stress (CMS) rat model. The CMS model mimics several psychopathological dimensions of depression ^17^, and adult SD rats that were exposed to the CMS regime exhibited obviously decreased sucrose preference and increased immobility time in the forced swim test (FST) (**Fig. 1E and 1F**). Notably, the concentrations of MGO were lower in the PFC and HC, two critical regions for stress response ^18, 19^, of CMS rats when compared with control rats (**Fig. 1A**), suggesting an MGO deficiency in the brains of rats after exposure to CMS. Accordingly, the plasma MGO levels also dropped remarkedly in CMS rats compared to vehicle (**Fig. 1A**). To determine whether reduced concentrations of MGO in CMS rats are associated with GLO1 activity, we employed quantitative real-time reverse transcription PCR (qRT-PCR) and immunoblot analysis. We found that both GLO1 transcription and expression are upregulated in the PFC and HC of CMS rats (**Fig. 1B** and **1C**). Together, these results suggest that reduced MGO levels in the brain underlie chronic stress-induced depressive behaviors.

**Figure 1.**
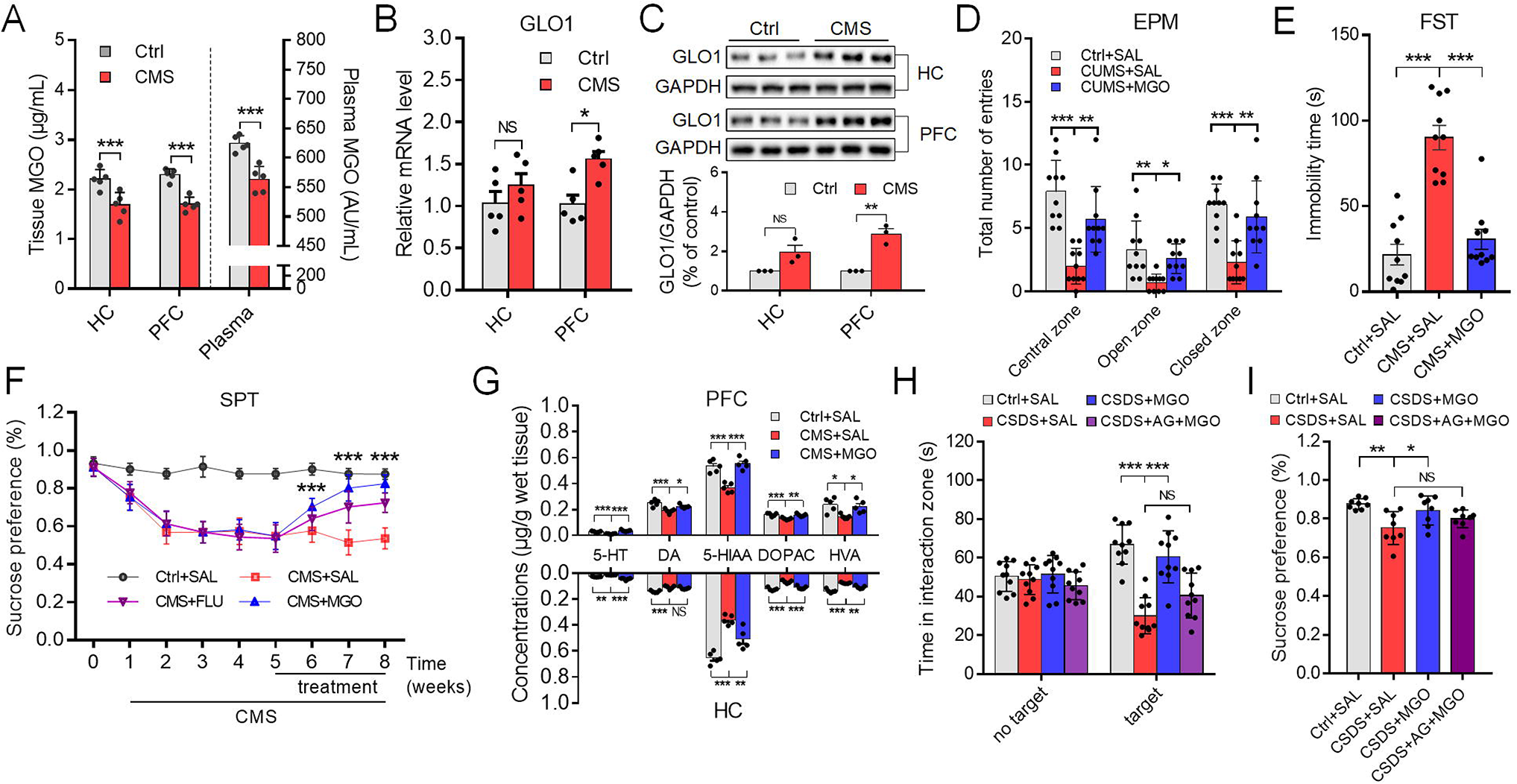
MGO increases resilience to chronic stress and restores neurotransmitter levels. (***A***) MGO concentrations in the HC, PFC and plasma of normal control rats (Ctrl) and CMS rats. MGO levels were measured by MGO-H1 protein adducts using ELISA assays (n = 5 rats per group). (***B***) Measurement of mRNA expression levels of GLO1 and BDNF by qRT-PCR in in the HC and PFC of normal rats (control) and CMS rats (n = rats 5 per group). (***C***) Quantification of GLO1 protein levels (percentage of protein to GAPDH) in the HC and PFC of normal rats (control) and CMS rats (n = 5 rats per group). (***D***) Comparison of immobility time in the forced swim test (FST) of normal rats (control) and CMS rats treated with either vehicle or MGO 5 mM/kg intraperitoneal (ip) per day for 3 weeks. MGO treatment significantly decreased the immobility time compared to CMS rats (n = 10 rats per group). (***E***) In the elevated plus-maze (EPM) test, CMS rats exhibited decreased number of entries to open arms, whereas treating with MGO 5 mM/kg ip per day for 3 weeks significantly increased the entry times to open arms (n = 10 rats per group). (***F***) In the sucrose preference test, in week 7, the CMS rats were treated with saline, MGO or fluoxetine (FLU) (n = 10 rats per group). (***G***) LC-MS/MS analysis of the concentrations of monoamines and their metabolites in the PFC and HC homogenates in rats (n = 5 rats per group). 5-HT, serotonin; DA, dopamine; 5-HIAA, 5-hydroxyindoleacetic acid; DOPAC, 3,4-dihydroxyphenylacetic acid; HVA, homovanillic acid. (***H***) Interaction zone times for the control mice and mice treated with saline, MGO or AG+MGO (i.c.v.) in conditions of no target and CD1 target (n = 10 rats per group). (***I***) The sucrose preference test for the mice after 7-d treatment with saline, MGO or AG+MGO (i.p.) (n = 10 rats per group). Data in A, B and C are analyzed using multiplet-test with Holm–Sidak correction. Data in D, E and G are analyzed using one-way ANOVA with Dunnett’s multiple comparisons test. Data in F, H and I are analyzed using two-way ANOVA with Sidak’s multiple comparisons test. Data are presented as mean ±s.e.m., ^*^P < 0.05, ^**^P < 0.01, ^***^P < 0.001, NS = not significant.

To test whether MGO could induce an antidepressant-like effect, we employed the FST and elevated plus-maze (EPM) test. The treatment of low-dose MGO (5 mM/kg per day) for 3 weeks significantly reduced the immobility time of CMS rats in the FST (**Fig. 1E**) and robustly increased their time in the open arms of the EPM (**Fig. 1D** and **Supplementary Fig. S1B**), indicating that MGO could have antidepressant-like effects. Of note, we observed that a 7-d treatment with MGO was sufficient to remarkedly increase the sucrose preference in rats with depressive behaviors (**Fig. 1F**). In contrast, a 3-week treatment with the typical antidepressant fluoxetine was necessary to increase the sucrose preference of CMS rats to the baseline levels, and a 1-week treatment with fluoxetine was ineffective. These results suggest that MGO may be a faster-acting antidepressant (**Fig. 1F**). Consistent with its antidepressant activity, MGO also greatly increased the levels of monoamine neurotransmitters such as serotonin and dopamine in both PFC and HC regions of CMS rats (**Fig. 1G**).

To further investigate whether MGO also promotes stress resilience in mice with depressive-like behavior, we employed another widely used stress model: the chronic social defeat stress (CSDS) paradigm (**Supplementary Fig. S1D**). Adult C57BL/6J mice that were subjected to CSDS were separated into susceptible and resilient subpopulations (**Supplementary Fig. S1E**). A physiological concentration of MGO (10 μM, lateral intracerebroventricular (i.c.v.) infusion) decreased the total duration of immobility in adult C57BL6/J mice. Notably, a preinfusion of aminoguanidine, a scavenger of free MGO, blocked MGO’s antidepressant-like effects (**Fig. 1H** and **1I**). We then considered whether MGO could reverse CSDS-induced social avoidance, which is a model of stress-induced psychopathology in humans ^20, 21^. After a 10-d CSDS protocol, the mice that were treated with vehicle exhibited an approximately 70% reduction in time spent in the interaction zone. In contrast, a 7-d treatment with MGO reversed the defeat-related behaviors in mice after CSDS (**Fig. 1H**), suggesting that MGO promotes resilience to chronic stress in adult mice.

### MGO promotes the synaptic plasticity in PFC and HC of CMS rats

To characterize the neurobiological mechanisms that underlie the antidepressant-like effects of MGO, we performed RNA-seq transcriptome profiling of the PFC and HC regions of three groups of rats, i.e., normal control rats and CMS rats treated with either saline or 5 mM/kg MGO for 3 weeks. Differential expression analysis showed that 289 genes in PFC and 278 genes in HC were specifically up- or downregulated in the MGO treatment group compared to the saline group (**Fig. 2A**), of which 63 genes are overlapped between these two regions (**Fig. 2A**). Notably, we found that MGO reversed the expression levels of 25.5% and 47.3% of the misregulated genes in PFC and HC of CMS rats to normal state. Among the top dysregulated genes were those encoding signaling molecules that are essential for differentiation and maintenance of adult neural stem cells in hippocampus, such as adenomatous polyposis coli (*Apc*) and BTG anti-proliferation factor 1 (*Btg1*); and important synaptic proteins such as the synaptotagmin (SYT) family, *Syt2*, *Syt11*, *Syt13*, and the disintegrin and metalloproteinase domain-containing protein 22 (*Adam22*) involved in neurogenesis (**Fig. 2A**). Gene ontology (GO) analysis of these differentially expressed genes revealed an enrichment of neuron-specific GO cellular component terms in both PFC and HC regions, including axon, dendritic spine, synapse, and also biological process terms such as calmodulin binding and calcium-mediated signaling, which are crucial for neurotransmitter release and neuro projection (**Fig. 2B**). As disruption of synapse plasticity is a major phenomenon in depression patients ^22^, these results suggest that MGO’s antidepressant effect is associated with the increased synaptic plasticity in PFC and HC of CMS rats.

**Figure 2.**
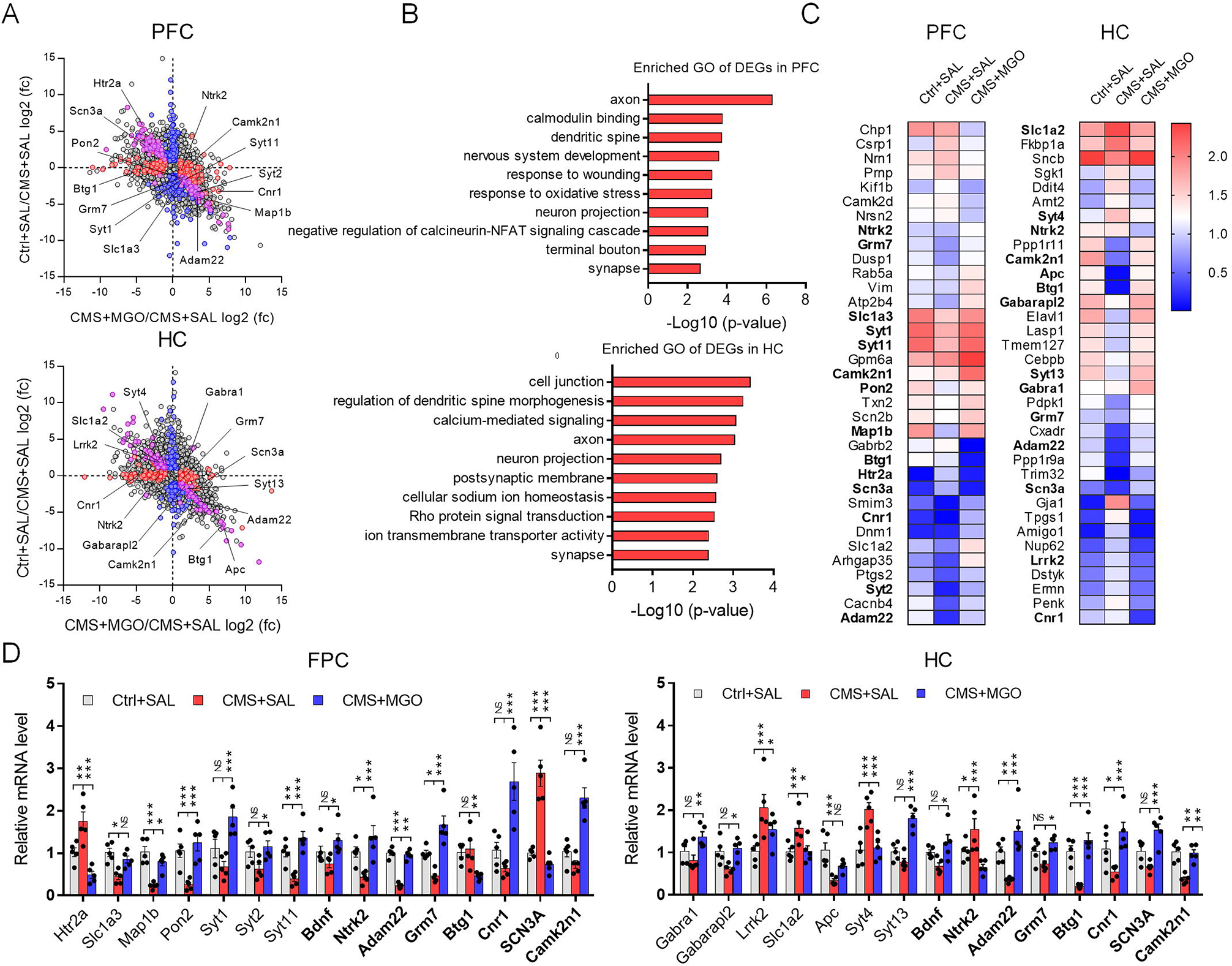
MGO promotes the synaptic plasticity in PFC and HC of CMS rats. (***A***) Scatter plot showing the differentially expressed genes (DEGs) in HC and PFC among the control rats and CMS rats treated with either vehicle or MGO 5 mM/kg intraperitoneal (ip) per day for 3 weeks. The horizontal axis represents the log_2_(fold change) of genes between CMS+MGO and CMS+SAL groups, and the vertical axis represents the log_2_(fold change) of genes between Ctrl+SAL and CMS+SAL groups. The colored dots represent DEGs with Llog_2_(fold change) ∟ > ∟ 2 and FDR < 0.05 (red, CMS+MGO vs CMS+SAL; blue, Ctrl+SAL vs CMS+SAL; pink, overlapped genes). (***B***) The enriched GO terms of MGO-induced DEGs in PFC and HC regions of CMS rats. (***C***) Heatmap showing the DEGs in the PFC and HC of different groups of rats. (***D***) qRT-PCR analysis of the DEGS related to synaptic plasticity in the PFC and HC lysates of different group of rats. Data are analyzed using two-way ANOVA with Sidak’s multiple comparisons test. Data are presented as mean ± s.e.m., n = 5 rats per group, ^*^P < 0.05, ^**^P < 0.01, ^***^P < 0.001, NS = not significant.

To validate the RNA-seq results, we focused on the genes associated with synaptic plasticity and neurogenesis, using qRT-PCR and immunofluorescence analyses (**Fig. 2D** and **Supplementary Fig. 1F**). The results showed that MGO’s regulation on the expression of these genes is in accordance with RNA-seq results. Of note, gamma-aminobutyric acid type A receptor related genes (*Gabra1*, *Gabarapl2*) in HC, which were previously implicated in anxious and depression ^15, 16^ were upregulated by MGO compared to CMS group. 5-hydroxytryptamine receptor 2A (*Htr2a*), mutations of which are associated with response to the antidepressant citalopram in MDD patients ^23^, is downregulated by MGO in the PFC region. In addition, MGO also significantly up-regulated the mRNA expression levels of *Bdnf* and *Adam22* that are involved in the positive regulation of neurogenesis (**Fig. 2D**). To test whether MGO modulates hippocampal neurogenesis *in vivo*, we carried out immunofluorescence analysis on the HC slices of CMS rats treated with MGO. The density of both BrdU+ new born neurons and BrdU+/NeuN+ neurons was decreased in the hippocampal dentate gyrus (DG) of CMS rats (**Supplementary Fig. 1F**). Whereas ip administration of MGO significantly increased the proportion of these cells (**Supplementary Fig. 1F**), indicating that MGO is capable of promoting hippocampal neurogenesis in CMS rats. Together these data suggest that MGO promotes the synaptic plasticity in PFC and hippocampal neurogenesis in CMS rats, and thus presents antidepressant-like effects.

### MGO activates TrkB signaling pathways and induces the expression of BDNF in PFC and HC of CMS rats

The neurotrophic factor BDNF and its receptor TrkB play an important role in regulating synaptic plasticity and hippocampal neurogenesis ^24, 25^. We thus examined whether MGO’s antidepressant effects is associated with the BDNF/TrkB signaling pathways in the brains of CMS rats. We found that the phosphorylation levels of two TrkB phosphorylation sites (Y515 and Y706) were significantly upregulated in MGO treatment group compared to the saline group. Accordingly, we also observed increased levels of downstream p-Akt, p-ERK1/2 and p-CREB in both HC and PFC areas of MGO-treated CMS rats (**Fig. 3A** and **Supplementary Fig. S2A**), which are important signals that regulate the neuronal survival and local axon growth ^26^. More importantly, MGO significantly increased the expression and release of BDNF in both of these brain areas (**Fig. 3A** and **Supplementary Fig. S2A**). To further validate MGO’s activation effects on TrkB signaling pathways *in vivo*, we adopted a single acute treatment of MGO (50 mM/kg) to CMS rats. The results demonstrated that MGO activates the TrkB signaling pathways in a time-dependent manner and maintains a persistent activation state in 9 h in both PFC and HC regions (**Fig. 3A and Supplementary Fig. S2B**). Together, these results demonstrate that MGO activates the BDNF/TrkB signaling pathways in both PFC and HC areas of CMS rats.

**Figure 3.**
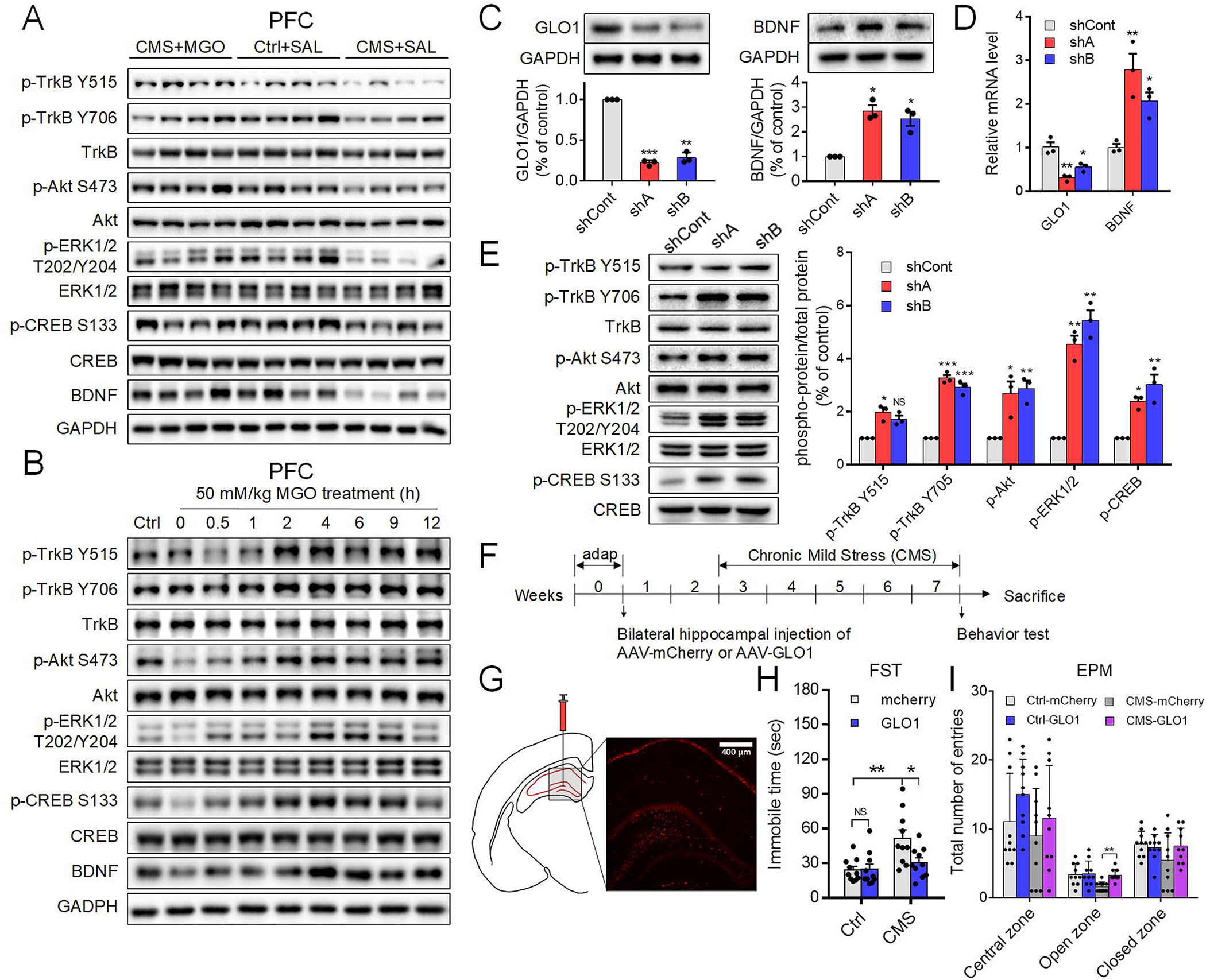
MGO activates TrkB signaling pathways and induces the expression of BDNF in PFC and HC of CMS rats. (***A***) Western blotting analysis of BDNF and other proteins involved in TrkB signaling pathway of PFC lysates of different groups of rats. Compared to CMS rats, MGO-treated rats (5 mM/kg, ip per day for 3 weeks) exhibited increased p-TrkB (Y706), p-Akt (S473), p-ERK1/2 (T202/Y204), and p-CREB (S133) immunoreactivities (n = 4 rats per group). (***B***) Representative western blot of PFC lysates of rats euthanized at various time points after ip administration of single dose MGO (50 mM/kg). MGO exhibits time-dependent activation on TrkB signaling pathways and BDNF expression. (***C***) Immunoblot analysis of protein expressions of GLO1 and BDNF (percentage of protein to GAPDH) in PC12 cells after shRNA-mediated GLO1 knockdown (shA and shB) compared to shCont. Representative western blots are shown on top of the quantitative plots. shCont, a non-targeting control shRNA; shA and shB are the shRNAs targeting GLO1 (n = 3 samples collected independently). (***D***) Measurement of mRNA expression levels of GLO1 and BDNF by qRT-PCR in PC12 cells after shRNA-mediated GLO1 knockdown compared to shCont (n = 3 samples collected independently). (***E***) Immunoblot analysis and quantification of protein expression, which highlights the increase of p-TrkB, p-Akt, p-ERK1/2 and p-CREB (percentage of phosphorylated protein to total protein) after shRNA-mediated GLO1 knockdown (n = 3 samples collected independently). (***F***) AAV injection and CMS experiment procedures for rats. On Week 7, the EPM and FST were conducted. (***G***) The injection site of AAV vector encoding GLO1 shRNA into the dentate gyrus (DG) region of HC (gray box, coronal section at +1.78 mm bregma), with an image of mCherry expression in the DG 3 weeks after injection (scale bar, 400 μm). (***H***) Comparison of immobility time in the forced swim test (FST) of control rats and CMS rats with GLO1-konckdown. GLO1-knockdown rats exhibited decreased immobility time compared to control CMS rats (n = 10 mice per group). (***I***) The elevated plus-maze (EPM) test showing that CMS rats with GLO1-knockdown exhibited significantly increased entry times to open arms compared to control CMS rats (n = 10 mice per group). Data in C, D and E are analyzed using one-way ANOVA with Dunnett’s multiple comparisons test. Data in D, E and G are analyzed using one-way ANOVA with Dunnett’s multiple comparisons test. Data in H and I are analyzed using two-way ANOVA with Sidak’s multiple comparisons test. Data are presented as mean ±s.e.m., ^*^P < 0.05, ^**^P < 0.01, ^***^P < 0.001, NS = not significant.

We then examined the concentration dependence and time course of MGO-mediated activation of TrkB signaling pathways in cultured PC12 cell lines *in vitro*. Incubation of MGO with PC12 cells for 1 h produced a concentration-dependent activation of the TrkB signaling pathway (**Supplementary Fig. S3A**). Consistently, MGO also increased the expression of BDNF mRNA in a dose-dependent manner (**Supplementary Fig. S3D**). Furthermore, incubation of MGO at 250 μM for 15 or 30 min both robustly increased the levels of p-TrkB, p-Akt, p-ERK1/2 and p-CREB (**Supplementary Fig. S3B**), which demonstrates that MGO provokes the activation of BDNF/TrkB signaling in a fast way. Importantly, the nuclear entry of p-CREB was also significantly promoted by 250 μM MGO (**Fig. 4C**) and the mRNA levels of BDNF robustly increased at 3 h, peaked at 12 h and was detectable until 24 h, which is consistent to its changes at the protein level (**Supplementary Fig. S3C and S3D**). Besides, the phosphorylation of TrkB and the downstream p-ERK1/2 and p-CREB stimulated by 250 μM MGO can last up to 12 h (**Supplementary Fig. S3C**). This long-lasting BDNF/TrkB signaling induced by MGO may enhance the synaptic transmission in adult hippocampus, and thus lead to antidepressant-like effects ^27, 28^.

**Figure 4.**
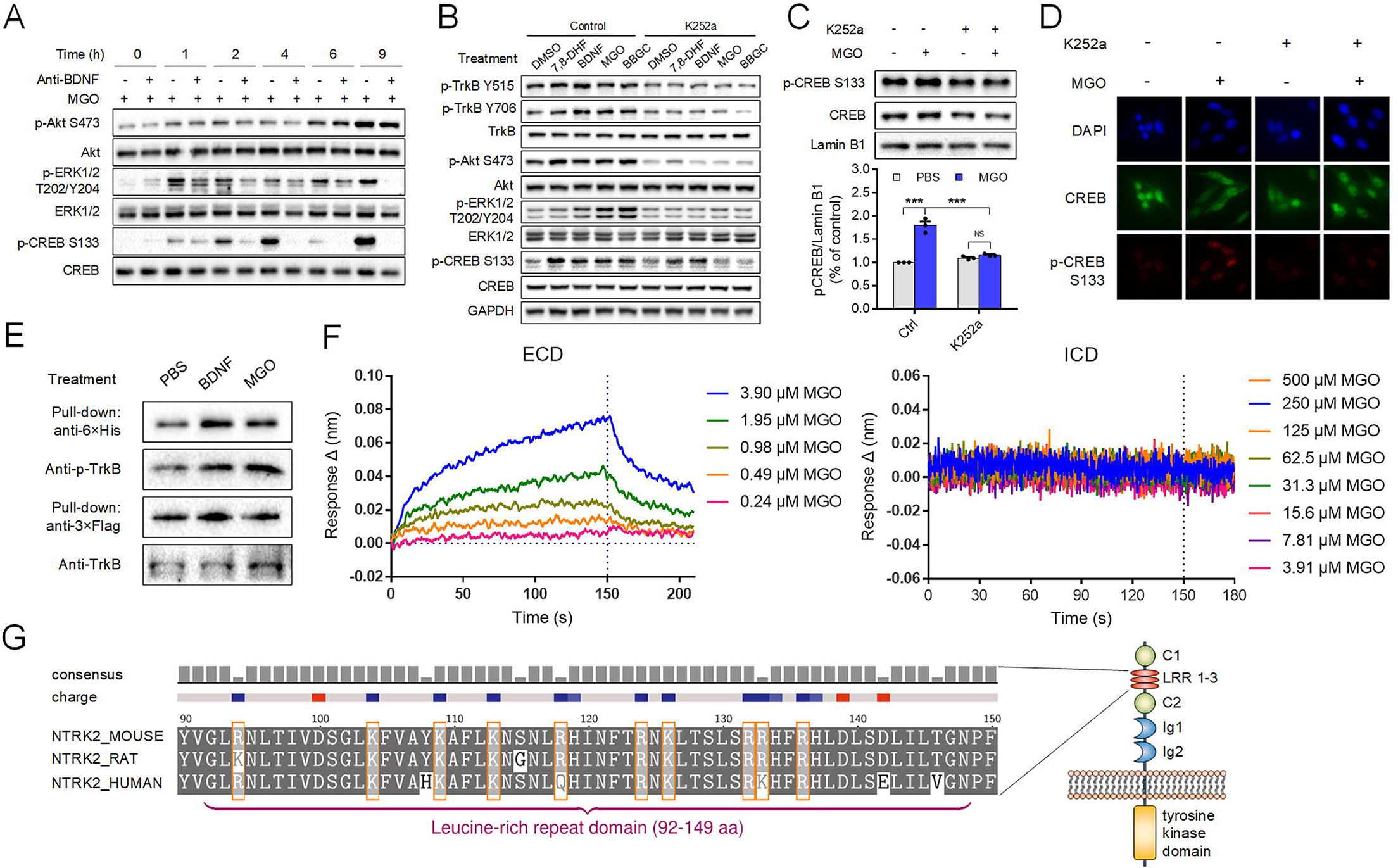
MGO directly binds to the extracellular domain of TrkB and provokes its dimerization and autophosphorylation. (***A***) Western blots of p-Akt (S473), Akt, p-ERK1/2 (T202/Y204), ERK1/2, p-CREB (S133) and CREB are shown. Either 250 μM MGO or additional 20 ng/mL anti-BDNF were preincubated for various periods of time with cultured PC12 cells. (***B***) Preincubating PC12 cells with 10 μM 7,8-DHF, 100 ng/mL BDNF, 250 μM MGO or 10 μM BBGC for 1 h significantly activated the TrkB signaling compared to that with 1% DMSO. While addition of 100 nM K252a markedly reduced the activation effects. (***C***) Treatment with 250 μM MGO for 1 h promoted the entry of p-CREB to nucleus, while the preincubation of PC12 cells with K252a for 30 min blocked this effect. Representative western blots of nucleus CREB and p-CREB (top) and the quantification of ratios of immunoreactivity of p-CREB (S133) to Lamin B1 (bottom) is shown. Data are analyzed using two-way ANOVA with Sidak’s multiple comparisons test. Data are presented as mean ±s.e.m., n = 3 samples collected independently, ^***^P < 0.001, NS = not significant. (***D***) Immunocytochemistry (ICC) assay depicting the total CREB (green) and the phosphorylation levels of CREB (red) in PC12 cells. Treatment with 250 μM MGO for 1 h induced the phosphorylation of CREB, while preincubating the PC12 cells with K252a for 30 min blocked this effect. DAPI (blue) was employed to stain the nuclei. (***E***) Flag-tagged and His-tagged TrkBs were co-transfected into HEK-293 cells. The cells were then treated with either PBS, 250 μM MGO or 100 ng/mL BDNF for 30 min. Subsequently, Flag-tagged TrkB was pulled down and monitored by 6×His-tagged antibody. (***F***) Biolayer interferometry (BLI) data depicting the association and dissociation sensograms at different concentrations of MGO for the interaction analysis of MGO with the extracellular and intracellular domain (ECD and ICD) of TrkB. (***G***) The enrichment of arginine (R) and lysine (K) in the third leucine-rich motif of TrkB from human, rat and mouse.

To investigate whether modulation of endogenous MGO levels could also regulate TrkB signaling pathways, two independent, non-overlapping short hairpin RNA (shRNA) lentiviral constructs were designed to knock down GLO1 in PC12 cells (**Supplementary Table S1**). The levels of GLO1 protein and its mRNA were significantly down-regulated, along with a significant increase in MGO concentrations in both of the shRNA groups (**Figs. 3C, 3D** and **Supplementary Figs. S2C**). We then tested whether this knockdown impacts the signaling transduction of BDNF/TrkB pathway. The results show that the expression levels of BDNF protein and its mRNA were both obviously enhanced (**Figs. 3*C* and 3D**). Correspondingly, silencing of GLO1 increased the levels of p-TrkB, p-Akt, p-ERK1/2 and p-CREB (**Fig. 3E**), indicating the activation of Akt and ERK signaling pathways. Additionally, the pharmacological inhibition of GLO1 by 10 μM S-p-bromobenzylglutathione cyclopentyl diester (BBGC) in WT PC12 cells also resulted in significantly increased MGO levels and enhanced TrkB signaling (**Supplementary Figs. S3E and 3F**). Reciprocally, the overexpression of GLO1 in PC12 cells significantly increased both the protein and mRNA levels of GLO1, along with decreased MGO levels (**Supplementary Figs. S2C, S2D and *S2E***). And this led to a reduction of p-TrkB, p-Akt, p-ERK1/2 and p-CREB levels, as well as the protein and mRNA levels of BDNF (**Supplementary Figs. S2D, *S2*E and *S2*F**), indicating that GLO1 overexpression causes functional inhibition of BDNF/TrkB signaling. To further demonstrate that modulation of endogenous MGO could have antidepressant-like effects *in vivo*, we utilized the adeno-associated viral (AAV) vector-mediated knockdown of GLO1 in the HC of CMS rats (**Fig. 3F** and **G**). The results showed that GLO1 knockdown CMS rats exhibited a significantly decreased immobility time in FST (**Figs. 3H**) and increased entries into open arms in EPM compared to CMS rats with the control vector (**Figs. 3I** and **Supplementary Figs. S2F**). Taken together, these results indicate that both exogenous and endogenous MGO can active the TrkB signaling pathway and induce the expression of BDNF.

### MGO directly binds to the extracellular domain of TrkB and provokes its dimerization and autophosphorylation

Next, we explored the molecular mechanisms underlying the fast and sustained activation effect of MGO on TrkB signaling pathways. To exclude the activation effects due to the accumulation of BDNF, we added the BDNF antibody to the cell cultures and found that it only partly abrogated MGO-induced phosphorylation of Akt, ERK1/2 and CREB in the first 9 h (**Fig. 4A** and **Supplementary Fig. S4A**). This result suggests that the apparent signaling activation effect caused by MGO does not completely attribute to BDNF. Further, we utilized K252a, a potent selective inhibitor for Trk receptors ^29^, to investigate whether MGO’s effect is associated with the kinase activity of TrkB. K252a potently inhibited the phosphorylation of TrkB, Akt and ERK1/2 that was stimulated by BDNF or 7,8-dihydroxyflavone (7,8-DHF), a known small molecule agonist of TrkB ^30^. And the levels of p-TrkB, p-Akt and p-ERK1/2 induced by 250 μM exogenous MGO or 10 μM BBGC were also significantly decreased by this inhibitor (**Fig. 4B** and **Supplementary Fig. S4B**). Additionally, K252a also decreased the p-CREB levels and its nuclear entry induced by MGO (**Figs. 4C** and **4D**). Given that MGO is previously reported as a competitive partial agonist of GABA_A_R ^15^ and activation of GABA_A_R is often related to TrkB signaling ^31, 32^, we further determined whether MGO’s activation effect on TrkB signaling is associated with GABA_A_R. The results show that MGO actives TrkB signaling regardless of the presence of a selective GAB_A_R antagonist, (−)-bicuculline methochloride (**Supplementary Fig. S4C**), suggesting that MGO directly actives TrkB phosphorylation independent of GABA_A_R.

To examine how MGO regulates the kinase activity of TrkB, we tested whether MGO induces the homodimerization and phosphorylation of TrkB. The pull-down assays showed that treatment of 250 μM MGO or 100 ng/mL BDNF both significantly provoked the homodimerization and autophosphorylation of TrkB (**Fig. 4E**), which indicates that MGO physiologically mimics the functions of BDNF. The biolayer interferometry (BLI) assay showed that MGO directly binds to the extracellular domain (ECD, 32-429aa) of TrkB with a dissociation constant (*K_d_*) of 5.31 μM (**Fig. 4F**). However, MGO does not interact with the intracellular domain (ICD, 454-821aa) of TrkB (**Fig. 4F**). Under physiological condition, MGO binds and modifies arginine, lysine, and cysteine residues in proteins ^33^. To explore the modification sites of MGO on the extracellular domain of TrkB, we analyzed the amino acid sequence of the binding regions between TrkB and BDNF. The binding regions in TrkB receptor include amino acid residues 103–181 that contain the third leucine-rich motif and the CC-2 domain and the Ig2 domain (342–394) ^34^. We found a significant enrichment of arginine (R) and lysine (K) in the third leucine-rich motif, whereas no arginine (R) or lysine (K) in the Ig2 domain (**Fig. 4G**), suggesting MGO binds to the arginine or lysine residues within the outer binging domain between BDNF and TrkB. Taken in sum, MGO selectively binds to the ECD of TrkB and functions as its agonist, and thus induces a fast and sustained activation of BDNF/TrkB signaling.

### GLO1 is associated with TrkB signaling in MDD patients and serves as a potential drug target for depression

To test whether in MDD patients GLO1 is associated with TrkB signaling, we performed a differential coexpression analysis on a gene expression profiling of brain samples from 34 MDD patients and 55 normal individuals ^35^. The differentially coexpressed genes (DCGs) of six brain areas, i.e., the dorsolateral PFC (DLPFC), HC, were obtained using an improved weighted correlation network analysis (WGCNA) approach ^36^ (see Methods, **Supplementary Table S2**). Pathway enrichment analysis of the six DCG sets against KEGG database identified the ‘neurotrophin signaling pathway’ as one of the most significantly enriched pathways in the DLPFC, AnCg, HC and NAcc areas (**Supplementary Fig. S5B**). Then, by employing the gene set enrichment analysis (GSEA) algorithm, GLO1 was found significantly correlated with the BDNF/TrkB signaling pathway in the DLPFC, HC areas of these samples (**Fig. 5A and Supplementary Fig. S5C**), further validating that MGO levels may be involved in regulating the BDNF/TrkB signaling in the DLPFC, HC areas of MDD patients.

**Figure 5.**
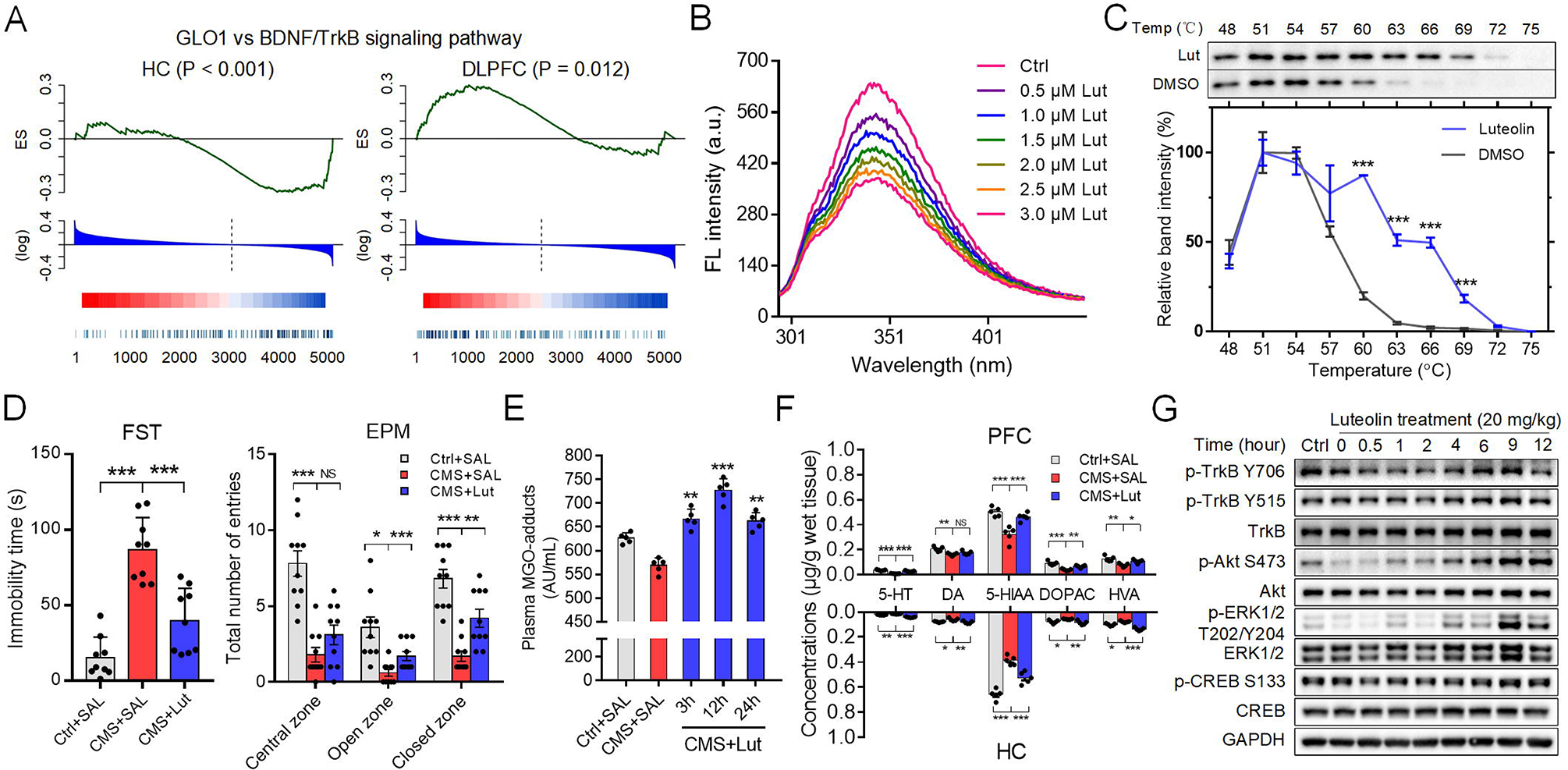
GLO1 is associated with TrkB signaling in MDD patients and serves as a potential drug target for depression. (***A***) GSEA analysis between GLO1 coexpression gene signatures and the BDNF/TrkB signaling pathway in HC and DLPFC areas of depression patients. HC, hippocampus; DLPFC, dorsolateral prefrontal cortex. (***B***) Fluorescence spectra analysis of GLO1 in the presence of various concentrations of luteolin. T = 298 K, λex = 280 nm, C_GLO1_ = 5 μM. (***C***) Representative western blot of CETSA in cell lysate for GLO1 targeted by 100 μM luteolin. Data are presented as mean ± s.e.m., n = 3 samples collected independently, ^*^P < 0.05, ^***^P < 0.001; two-way ANOVA with Sidak’s multiple comparisons test between luteolin treatment group and DMSO group. (***D***) Luteolin (10 mg/kg) significantly increased the number of entries of rats in central and open zones during the FST test and decreased immobility time in the EPM test compared to vehicle treatment, indicating an antidepressant-like response (n = 10 rats per group). (***E***) Plasma MGO levels of CMS rats after treated with luteolin for 3, 12, 24 h (n = 5 rats per group). (***F***) Measurements of monoamines and their metabolites levels in the PFC and HC of different groups of rats (n = 10 rats per group). 5-HT, serotonin; DA, dopamine; 5-HIAA, 5-hydroxyindoleacetic acid; DOPAC, 3, 4-dihydroxyphenylacetic acid; HVA, homovanillic acid. (***G***) Time course of TrkB, Akt, ERK1/2 and CREB and their phosphorylation levels as assessed by western blot in the HC of CMS rats after single dose treatment with luteolin (20 mg/kg). Data in D, E and F are analyzed using one-way ANOVA with Dunnett’s multiple comparisons test. Data in H and I are analyzed using two-way ANOVA with Sidak’s multiple comparisons test. Data are presented as mean ±s.e.m., ^*^P < 0.05, ^**^P < 0.01, ^***^P < 0.001, NS = not significant.

The antidepressant effects of MGO also makes GLO1 a potential therapeutic target for depression. By application of two in house systems pharmacology-based drug targeting tools, i.e., WES ^37^ and TCMSP ^38^, a flavone molecule, luteolin, was screened out as a candidate compound that may target GLO1 (**Supplementary Fig. 6A**). To validate the interaction between luteolin and GLO1, we employed both in vitro fluorescence spectroscopy and in vivo cellular thermal shift assay (CETSA) experiments ^39^. The fluorescence spectroscopy results showed a *K_d_* value of 47.17 μM for luteolin binding to GLO1 (**Fig. 5B**). And the *in vivo* CETSA also validates their binding in intact cells (**Fig. 5C**) with the EC_50_ concentration of 45.23 μM for luteolin at which half-maximal thermal stabilization of GLO1 in PC12 cells was observed (**Supplementary Fig. S6E**).

To explore the binding mode of luteolin with GLO1, we performed molecular docking coupled with molecular dynamic (MD) simulations. Three residues, E99, Q33 and F62 were identified as crucial residues for the binding, as that in GLO1 active pocket they formed hydrogen bonds and an edge-to-face aromatic interaction with luteolin (**Supplementary Fig. S6C**). Regarding luteolin, its carbonyl, resorcinol and pyrocatechol substituents form H-bonds and hydrophobic-interactions with GLO1, demonstrating the key roles of these groups in luteolin’s binding within the active site of GLO1 (**Supplementary Fig. S6C**). More importantly, we found that the hydroxyl group of the resorcinol is freely buried into the positively charged mouth, a big hydrophobic tunnel constituted by residues F62, F67, L69 and T101 (**Supplementary Fig. S6C** and **S6D**). To further investigate whether this hydrophobic cavity is beneficial for the binding, we modified the hydroxyl group of the resorcinol into a bulkier group, i.e., 1, 2-dimethoxyethane, and obtained a new derivative, named as lutD (**Supplementary Fig. S6A** and **S7**). Intriguingly, fluorescence spectroscopy analysis presents a 1.5-fold increase of the association constant (*K*_a_) (from 2.12 × 10^4^ L•mol^−1^ to 3.22 × 10^4^ L•mol^−1^) for lutD compared with luteolin (**Supplementary Fig. S6E**), indicating that the bulky groups in resorcinol contribute to a more potent GLO1 binding. Actually, MD simulations on lutD and GLO1 shows that this mouth zone is empty and thus provides sufficient space for bulky substituents (**Supplementary Figs. S6C and S6D**). Besides, the hydrophobic interaction between the methoxyethane group of lutD and F67, L160 (**Supplementary Fig. S6D**) also strengthens the interactions between lutD and GLO1. All these results demonstrate that luteolin is a potent binder of GLO1, in which H-bonds and hydrophobic interactions are important for the binding.

We next evaluated whether luteolin exhibits antidepressant effects on CMS rats. The results show that ip administration of 10 mg/kg luteolin per day for 3 weeks markedly reduced the immobility time of rats in FST and increased their number of entries in the open arms in EPM (**Fig. 5D** and **Supplementary Fig. S6F**). We also found that chronic luteolin treatment increased the plasma MGO levels and the concentrations of neurotransmitters in HC and PFC areas of rats (**Figs. 5E** and **5F**). These results demonstrate that luteolin exerts antidepressant effects on CMS rats by targeting GLO1. Besides, similar to MGO, luteolin treatment also promoted hippocampal neurogenesis as indicated by obviously increased number of BrdU+ or BrdU+/NeuN+ cells in the hippocampal DG of CMS rats (**Supplementary Fig. S6G**). Additionally, single dose treatment of 20 mg/kg luteolin potently activated the BDNF/TrkB signaling in a time-dependent manner in the HC of CMS rats (**Fig. 5G**). These data reveal that through targeting GLO1, luteolin increases MGO concentrations in CMS rats and subsequently triggers the BDNF/TrkB signaling pathway, resulting in desirable antidepressant effects.

## Discussion

In this study, we demonstrated that MGO functions as an endogenous agonist of TrkB. We discovered that MGO directly and selectively binds to the ECD of TrkB and stimulates its dimerization and autophosphorylation. Analysis of amino residues of the binding region between BDNF and TrkB revealed that there is a significant enrichment of arginine and lysine in the third leucine rich motif, suggesting potential MGO modification sites on TrkB. Both in vitro and in vivo experiments showed that MGO triggers a rapid and sustained activation of TrkB-mediated Akt/CREB signaling, which rapidly increases the expression of BDNF, thus being beneficial to MDD. Clinically, increasing the BDNF level in brain is of particular therapeutic interests for depression ^40, 41^. However, current clinical trials using recombinant BDNF in patients always turn out disappointing due to series of reasons like poor delivery, short half-life, and potential side effects ^42^. Our finding that MGO at normal concentrations acts as a switch on activating the BDNF-feedback loop provides a new feasible way to increase the BDNF levels for MDD patients.

Importantly, we found that MGO levels in the PFC and HC are significantly decreased in the CMS rats, whereas treatment of low-dose MGO (5 mM/kg) per day for 3 weeks reverses the MGO levels back to normal concentrations and exhibits desirable antidepressant effects. MGO also promotes the synaptic plasticity and neurogenesis of PFC and HC neurons. Besides, we found that chronic administration of MGO does not cause any weight gains in rats (**Supplementary Fig. S1C**), avoiding the major side effects produced by current clinically-used tricyclic antidepressants and monoamine oxidase inhibitors ^43, 44^. Except for depression, some other psychiatric and neurological disorders such as the epilepsy and Parkinson’s disease also possess several common pathological features, like synaptic density reduction and neuron atrophy ^45, 46^. Thus, manipulating endogenous MGO concentrations in brain provides a new alternative way for treating both MDD and these diseases with more favorable tolerability and efficacy.

The present study also highlights the potential of GLO1 as a depression target (**Fig. 6**). And based on this, luteolin was screened out as a GLO1 inhibitor and its binding affinity as well as the interaction features with GLO1 were deeply explored. As a natural flavonoid, luteolin exists in many plants such as *Apium graveolens*, *Petroselium crispum*, and *Capsicum annuum*, with antioxidant, anticancer, memory-improving, and anxiolytic activities ^47^. In the present study, chronic administration of luteolin for 3 weeks significantly improves the depression-like behaviors in CMS rats. Luteolin also increased the number of new-born neurons in the HC of CMS rats (**Supplementary Fig. S6G**), which provides a reasonable explanation of why luteolin has strong neuroprotective effects both *in vitro* and *in vivo* ^47, 48^. Importantly, long-term treatment of luteolin does not exhibit any apparent toxicity in rats (10 mg/kg, i.p. for 3 weeks), which corresponds well with the intraperitoneal LD50 of luteolin in rats (411 mg/kg) ^49^. In conclusion, our work not only presents a novel mechanism for MGO’s effect on promoting stress resilience, but also identifies a natural product luteolin as a promising antidepressant.

**Figure 6.**
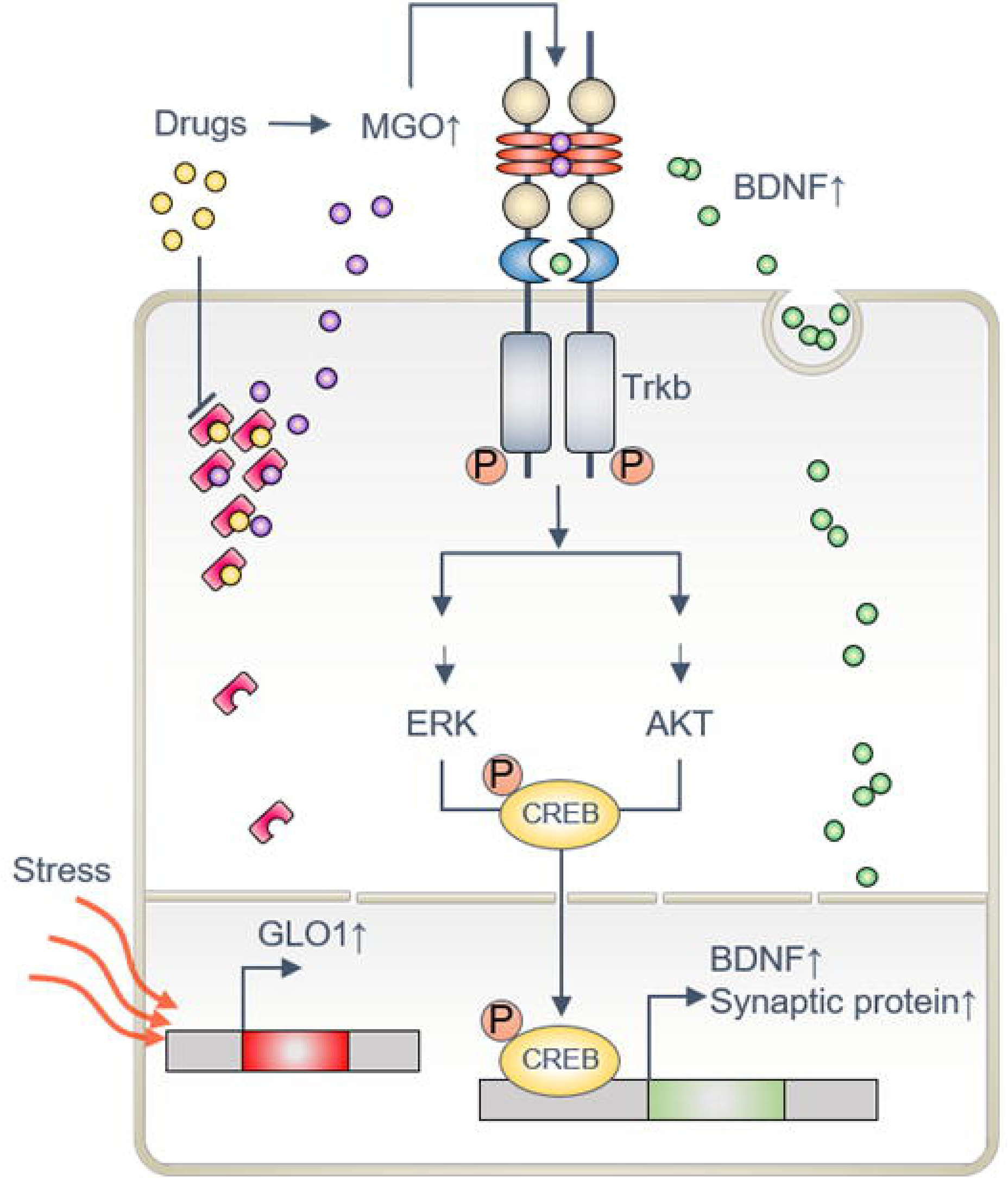
The molecular mechanisms underlying the antidepressant effects of MGO. Under normal conditions, MGO binds to the extracellular domain of TrkB and provokes its dimerization and autophosphorylation, which then activates the downstream Akt and ERK signaling, as well as the phosphorylation of the transcription factor CREB. This effectively induces the expression of BDNF and forms a BDNF-positive feedback loop, which further promotes the synaptic plasticity of nerons. Whereas under depressive state, the concentrations of MGO are significantly decreased partly due to the increased expression levels or enzyme activities of GLO1. This results in inactivation of the BDNF/TrkB signaling. GLO1 inhibitors, such as luteolin, can effectively increase the concentrations of MGO and thus exert antidepressant effects.

## Materials and methods

### Animals

Adult male C57BL/6J mice (7–9 week old, 22–24 g) and adult Sprague-Dawley (SD) male rats (180 - 200 g) were purchased from Vital River Laboratory Animal Technology Co., Ltd. (Beijing, China). Three to four mice were housed in a plastic cage (300 × 170 × 120 mm) at 24 ± 1 °C. Both mice and rats were group-housed with standard laboratory bedding and conditions (12-h light/dark cycle, 22 ± 1°C, ad libitum access to food and water) for 1 week prior to the experiments. The experimental procedures performed in this study were in line with the Guidelines of NIH for the Use of Laboratory Animals as well as approved by the Northwest University Animal Care and Use Committees.

### Drugs

The following drugs were used in this study, including methylglyoxal solution (Sigma-Aldrich, M0252), S-p-bromobenzylglutathione cyclopentyl diester (Sigma-Aldrich, cat. no. SML1306), K252a (Abcam, cat. no. ab120419), luteolin (TCI chemicals, TT2682), 7,8-dihydroxyflavone (7,8-DHF) (Abcam, cat. no. ab120996), and recombinant human BDNF protein (R&D Systems, cat. no. 248-BD-025).

### CMS procedure

To induce the physical, behavioral, biochemical, and physiological phenotypes of depression, all rats were subjected to a schedule of mild psychosocial stressors for 8 weeks (**Supplementary Fig. S1A**). The stressors included alterations of the bedding (sawdust change, removal or damp of sawdust, substitution of sawdust with 21°C water, rat, or cat feces for 1-3 h), cage-tilting (45°,1-3 h) or shaking (5-20 min), cage exchange (rats were positioned in the empty cage of another male for 2-6 h), predator sounds (5-20 min), and alterations of the length and time of light/dark cycle. To increase stress intensity, stressors were combined after the first week of the CMS regimen ^50^. Physical changes were assessed weekly by measuring the body weight of the animal. Additionally, in order to evaluate the neurogenesis by immunocytochemistry, rats were ip injected with BrdU (2 × 150 mg/kg) at the day before treatments. All rats were randomly assigned to three experimental groups, i.e., non-stress control + saline (SAL); stress (CMS) + SAL; stress (CMS) + drugs (5 mM/kg/day of MGO or 10 mg/kg/day of luteolin). To avoid the possible bias induced by the behavioral tests, the rats in each group were divided into two sets (n = 10 per set). The first set provided tissue samples used for morphological analysis, western blotting and qRT-PCR analyses, while the second one was adopted for behavioral assessments. During the last 3 weeks of the CMS protocol, rats were injected ip daily with drugs.

### Behavioral assessment

At the end of the CMS protocol, the behavioral tests EPM and FST were conducted. Two days were given for rats between exposures to different behavioral assessments and all behavioral testing experiments were carried out during the daily light phase (09:00 am – 05:00 pm).

#### Elevated plus-maze test

Rats were placed in the center of a standard EPM apparatus (two open and two closed 33 cm × 5 cm arms) for examining the anxiety-like behaviors of rats. The exploratory activity was measured over a 5-min period. Then, the percentage of time spent in the open arms, an index of anxiety-like behavior, the number of entries in the closed arms, an indicator of locomotion, were determined.

#### Forced swim test

In brief, SD rats were placed in a glass cylinder filled with water (23°C; 30 cm deep) and a 5-min swim test session was video-recorded. Employing an automated video tracking system, the time spent immobile during the last 4 minutes of the test and the latency to immobility were scored. An increase in immobility time indicates a higher degree of depressive state. All tests were analyzed by TopScan (CleverSys, Inc., CSI) system.

#### Sucrose preference test

The sucrose preference test was conducted based on previously described procedures ^21^. Briefly, the mice or rats were habituated to a 1% sucrose solution for 2 days before 2 additional days of choice testing. The position of the water and sucrose bottles was switched every 12 h to ensure that the mice or rats did not develop a side preference. After this period, the mice or rats were deprived of food and water for 24 h, and exposed to two bottles filled with either 1% sucrose or water for 12 h in the dark. Total consumption of each fluid was measured and sucrose preference was calculated as a percentage (amount of sucrose consumed (bottle A) × 100 / total volume consumed (bottles A + B)). To avoid the bias related to body weight, this variable was calculated as an amount of consumed sucrose in mg per gram body weight × 100.

### CSDS procedure and the social interaction test

The CSDS protocol was conducted as previously described ^20^. Briefly, a singly C57BL/6J mouse was exposed to a different aggressive CD1 mouse for 5 min each day for a total of 10 days. Following 5 min of contact, the intruder mouse was housed across a perforated plastic divider that allowed visual, olfactory and auditory contacts, but prevented physical interaction from the aggressor CD1 mouse, for the remainder of the 24-h period. Non-defeated control mice were housed opposite to another C57BL/6 mouse and the controls were changed daily. Following the 10-day CSDS protocol, the avoidance behaviors were tested to separate the susceptible and unsusceptible subpopulations, the interaction ratio was then calculated as 100 × (time spent in the interaction zone with an “Target”)/(time spent in the interaction zone with “No Target”) and an interaction ratio of 100 was set as a cutoff. The mice with scores <100 were considered susceptible, and those with scores ≥100 were considered resilient. On day 13, each mouse was implanted with an osmotic minipumps (Alzet, model 1007D; 0.5μL/h, 7 d) with a reservoir containing either vehicle (aCSF) or MGO (10 μM) in the right lateral ventricle. To block MGO-induced effect, rats were pre-treated with AG (1 g/L) via drinking water for 48 h (on day 11) before pumps implantation. On day 20, test mice (defeated mice and non-defeated control mice) were subjected to the social interaction test. The approach-avoidance behavior of a test mouse to a CD1 mouse was recorded with a video tracking system. The social interaction test consisted of two 150 s sessions. In the first phase, ‘no target’ session, the mouse could explore freely in a square-shaped open-field arena (43.5 × 43.5 cm) possessing a wire-mesh cage (10 × 6.5 cm). For the second ‘target’ session, the mouse was reintroduced into this arena with an unfamiliar CD1 mouse placed into the wire mesh cage. The wire mesh cage allowed visual, olfactory and auditory interactions between the test mouse and the target CD1 mouse but prevented direct physical contact. After each trial, the apparatus was cleaned with a solution of 70% ethanol in water to remove olfactory cues and all the behavioral tests were conducted in a dark room. The duration of time that the mice spent in the interaction zone and corner zone was obtained using TopScan (CleverSys, Inc., CSI) system.

### Quantification of MGO levels

Analysis of the levels of MGO in the plasma (Hycult Biotech, HIT503) and cell lysis (Cell Biolabs, STA-811) was carried out using specific enzyme-linked immunosorbent assays (ELISA) following the manufacturer’s instructions. Plasma was collected from aortaventralis and separated into Eppendorf tubes that contained EDTA to inhibit coagulation effect. The mixed blood and EDTA were then inversed and placed on ice for 20 min. After the inversion, the mixture blood sample was centrifuged for 15 min at 1500×g at 4°C. The supernatant was then collected and transferred to a fresh polypropylene tube for analysis. To detect MGO-adducts in cell lysate, 50 μg of total protein from fresh tissues or cell cultures were prepared using a Qproteome Mammalian Protein Prep Kit (Qiagen, Germany) in accordance with the manufacturer’s protocol.

### Cell culture and transfection

PC12 and HEK293 cells were cultured in DMEM medium supplemented with 10% fetal bovine serum (FBS), 1% (v/v) penicillin and streptomycin at 37°C under an atmosphere containing 5% CO_2_. To examine the relationship between GLO1 and TrkB signaling, we modulated the expression levels of GLO1 by lentiviral (LV)-mediated overexpression or shRNA knockdown in PC12 cells. The pLV-CMV-MCS-3flag-EF1-zsgreen-puro vector and the pLV-U6-MCS-zsgreen-puro vector for GLO1 overexpression and GLO1 knockdown were generated and obtained from Sangon Biotech (Shanghai) Co., Ltd.. The sequences of the non-targeting shRNA or GLO1-targeting shRNAs were listed in **Supplementary Table S1**. At 24 h before transfection, approximately 5.0 × 10^5^ cells were seeded in 6-well cell culture dishes. On the next day, they were infected with recombinant lentivirus in the presence of polybrene (5 μg/mL). The medium was replaced 12 h post-transfection with DMEM supplemented with 10% FSB, and cells were allowed to recover for additional 5 days. After transfection, cells were harvested, lysed and then employed for further western blotting analysis. To examine the homodimerization of TrkB induced by MGO, HEK293 cells were co-transfected with 3×Flag-tagged TrkB and 6×His-tagged TrkB. After transfection, these cells were then incubated with either PBS, 250 μM MGO or 100 ng/mL BDNF for 1 h for pull-down assay by Dynabeads His-Tag Pulldown (Invitrogen, cat. no. 10104D) according to the manufacturer’s instructions. For *in vitro* mutant studies, HEK293 cells were transfected with human 3×Flag-tagged GLO1 wild type and mutants (Q33E, E99A, F62A). Using Anti-DYKDDDDK Affinity Resin (Invitogen, cat. no. A36803), the Flag-tagged GLO1 wild type and mutants were purified for subsequent experiments.

### Quantitative reverse transcription-PCR

Total RNA was prepared from fresh tissues or cell cultures using the RNeasy mini kit (Qiagen, Germany) according to the manufacturer’s protocol. cDNAs were synthesized from 100 ng of total RNA using PrimeScript™ RT reagent Kit with gDNA Eraser (Takara, cat. no. RR047). Subsequently, qRT-PCR was performed employing the StepOnePlus Real-Time PCR System (Applied Biosystems) with SYBR Premix Ex Taq II (Takara. cat. no. RR820). Using Primer 3 software, specific gene primers (Sangon Biotech) used for qRT-PCR assays were designed and their sequences are listed in **Supplementary Table S3**. The expression levels of these genes were normalized to that of beta-actin and the fold changes in gene expression were computed by setting the gene expression levels in control samples as one.

### Western blot analysis

The expression levels of related proteins were examined by western blot experiments. A total of 30 μg protein samples were firstly separated by sodium dodecyl sulfate polyacrylamide gel electrophoresis (SDS-PAGE) and subsequently transferred onto poly vinylidene fluoride (PVDF) membranes. After blockade with 5% skimmed milk in TBST at room temperature for 1 h, the membranes were probed with specific primary antibodies to p-TrkB Y706 (Abcam, cat. no. ab197072, 1:1000), TrkB (Abcam, cat. no. ab18987, 1:1000), p-Akt S473 (Cell Signaling, cat. no. 4058, 1:1000), Akt (Cell Signaling, cat. no. 92715, 1:1000), p-ERK1/2 T202/Y204 (Cell Signaling, cat. no. 9101, 1:1000), ERK1/2 (Cell Signaling, cat. no. 4695, 1:1000), p-CREB S133 (Abcam, cat. no. ab32096, 1:5000), CREB (Abcam, cat. no. ab178322, 1:500), BDNF (Abcam, cat. no. ab108319, 1:5000), GLO1 (Abcam, cat. no. ab96032, 1:2000), GAPDH (Cell Signaling, cat. no. 5174, 1:1000), Anti-DDDDK tag (Abcam, cat. no. ab49763, 1:1000), Anti-6X His tag (Abcam, cat. no. ab18184, 1:1000) and then incubated with species-specific peroxidase-conjugated secondary antibodies (antibody to rabbit, Abcam, cat. no., ab6721, 1:20000; antibody to mouse, Abcam, cat. no., ab6789, 1:10000). Using Immun-Star™ WesternC™ Chemiluminescence Kit (Bio-Rad, cat. no. 1705070), the protein bands were detected and the optical densities of the bands were quantified by Image-Pro Plus v6 (Media Cybernetics, Silver Spring, MD).

### Immunocytochemistry

Cultured PC12 cells were pretreated with either vehicle or 100 nM k252a for 30 min and then treated with 250 μM MGO for 1 h. Subsequently, the cells were fixed in 4% formaldehyde for 10 min and incubated with primary antibodies overnight at 4°C, followed by incubation with the secondary antibodies. The following antibodies are used: p-CREB S133 (Abcam, cat. no. ab32096, 1:250), CREB (Abcam, cat. no. ab178322, 1:500), Goat Anti-Mouse IgG H&L (Alexa Fluor^®^ 488) (Abcam, cat. no. ab150117, 1:1000), Goat Anti-Rabbit IgG H&L (Alexa Fluor^®^ 594) (Abcam, cat. no. ab150080, 1:1000). Images were obtained using a DMI 4000B microscope (Leica, Germany) with DFC310 FX camera (Leica, Germany) and Leica application suite, version 4.2.0 (Leica Microsystems).

### Immunohistochemistry

Animals were deeply anaesthetized with sodium pentobarbital (50 mg/kg, ip). Brains were then removed and stored at −80°C. Coronal cryosections (20 μm) of the entire hippocampus were prepared with a cryostat (Leica, Wetzlar, Germany). After section, the slides were immediately immersed into 4% paraformaldehyde for 15 min at 4°C. For BrdU labeling, sections were denatured by treatment with 1N HCl for 10 min at 4°C followed by 2N HCl for 20 min at 37°C and then neutralized in 0.1 M borate buffer (pH 8.5) for 10 min. After washed with 0.3% Triton X-100 in PBS, the sections were then incubated with blocking solution containing 10% donkey serum (Jackson ImmunoResearch, cat. no. 017000121) in PBS for 60 min. Afterward, the sections were incubated in the primary antibodies (anti-BrdU, Abcam, cat. no. ab1893, 1:100; anti-NeuN, Abcam, cat. no. ab177487, 1:500) for 16 h at 4°C. Finally, sections were washed three times with PBS and then incubated with the corresponding Alexa-labeled secondary antibodies (donkey anti-rabbit, Abcam, cat. no. ab150061, 1:500; donkey anti-sheep, Abcam, cat. no. ab150180, 1:500) for 1 h at room temperature. After staining, slides were photographed using Leica DM 4000B microscope (Leica, Germany) fitted with a digital camera DFC450 C (Leica, Germany), and Leica application suite, version 4.2.0 software (Leica Microsystems) was used for image analysis. For quantification of BrdU-labeled cells, a modified unbiased stereology protocol was used as described previously ^51^. Briefly, we collected 5 rats for each group. One in six series of sections of each brain were counted in the dentate gyrus throughout the hippocampus. Results were multiplied by ten to obtain the total number of BrdU-positive cells per dentate gyrus. The average of these values for the individual brain was used for statistical analysis.

### Quantification of neurotransmitters and their metabolites by LC-MS/MS

To determine the effects of drugs on the depression rats, we measured the levels of the neurotransmitters in HC and PFC areas of rats through a rapid-resolution liquid chromatography–mass spectrometry (LC-MS)/MS analysis. The tissues sample preparation for the LC-MS/MS analysis were obtained from a liquid-solid extraction approach. Using a Waters ACQUITY UPLC system (Waters Corp., Milford, MA, USA) coupled to a QTRAP 5500 mass spectrometer (Applied Biosystems, Foster City, CA, USA), all analyses were performed. The chromatographic separation was carried out on Acquity UPLC HSS T3 column (2.1 mm × 100 mm, 1.8 μm) and BEH Amide column (2.1 mm × 100 mm, 1.7 μm), and the mass spectrometric measurements were carried out on a QTRAP 5500 quadrupole time-of-flight mass spectrometer equipped with an electrospray ionization source (ESI) operating in the positive and negative ionization modes, which were used for detection (Agilent Technologies). For both platforms, results were denoted as data matrices, in which integrated peak areas were used for determining the metabolites in all the samples.

### RNA-seq

On Week 7, the hippocampi and PFC of the rat brains were dissected from one normal rat and two CMS rats treated with either saline or 100 mM/kg MGO. Total RNA was extracted and purified from ~30∟mg tissue using the RNeasy mini kit (Qiagen, Germany) according to the manufacturer’s protocol. RNA-seq was performed at Genminix Informatics Ltd., Co. (Shanghai, China) using Illumina HiSeq ×10 sequencing platform. The original sequencing data have been uploaded to the GEO database (GSE119482).

### Identification and function enrichment analysis of differentially expressed genes

Quality control of raw sequences from each sample was investigated using FastQC (v0.11.5). Reads were then mapped to the Ensembl rat genome sequence (Rnor_6.0) using HISAT2 (v2.0.4) ^52^ and assembled into transcripts by StringTie (v1.2.4) ^53^ using a UCSC rn6 GTF (general transfer format) file. FPKM (fragment per kilobase per million mapped reads) values were used to estimate the transcript abundance. The ballgown ^54^ R package was employed to detect differentially expressed genes (DEGs) between normal, CMS and MGO-treated CMS rats. The Gene Ontology (GO) and KEGG pathway enrichment analyses of DEGs were performed using the DAVID bioinformatics resources 6.8 ^55^.

### Biolayer interferometry

The kinetics of MGO binding to the extracellular or cytoplasmic domain of recombinant human TrkB protein was assessed using biolayer interferometry with an Octet K2 system (ForteBio). All of the experiments were performed at 30°C under buffer conditions of PBST (0.1% Tween 20), pH 7.4, 8% DMSO. NI-NTA biosensors (FortéBio Inc., Menlo Park, CA) were used to capture TrkB proteins onto the surface of the sensor. After reaching baseline, sensors were moved to association step for 60 s and then dissociated for 60 s. Curves were corrected by a double-referencing technique, using both NI-NTA pins dipped into the experimental wells and a buffer-only reference. After double referencing corrections, the subtracted binding interference data were applied to the calculations of binding constants using the FORTEBIO analysis software (Version: 9.0.0.10) provided with the instrument.

### Stereotactic injection of adeno-associated virus

Adeno-associated viral vectors (AAVs) serotype 2 expressing either mCherry or GLO1 were generated and purified by Sangon Biotech (Shanghai) Co., Ltd. The rats used for the stereotactic injection were weighted from 280 g to 320 g. These rats were deeply anaesthetized with isoflurane gas (3.0% isoflurane for induction and 2.0% for maintenance) and placed into a stereotactic apparatus (RWD Life Science Inc.). Stereotaxic surgery was performed to deliver viruses into the hippocampus of these rats. AAV-GLO1 vector or AAV-mCherry was bilaterally infused using stereotaxic coordinates according to the rat brain atlas ^56^. By using a precision Hamilton micro-syringe with a 26 G needle, a total of 2 μL viral solution was bilaterally injected into the hippocampus (AP −3.6 mm, ML ± 2.0 mm, DV 3.4 mm). Viruses were infused at a speed of 0.2 μL/min for 10 min and the needle was left in place for additional 10 min before slowly being withdrawn. The rats were recovered on hot pad (37°C) until waken up and returned to cages, and then they were allowed to recover for 3 weeks before starting the behavioral paradigm.

### Gene expression data of MDD patients

Normalized gene expression data for MDD patients were collected from the GEO database (GSE45642) ^35^. GSE45642 includes a total of 670 whole-genome gene expression profiles of six brain areas, i.e. DLPFC, AnCg, HC, AMY, NAcc and CB for 55 normal controls and 34 MDD patients.

### Differentially coexpressed genes

To identify the DCGs in the six brain areas of MDD patients, we applied an I-WGCNA method ^57^ (**Supplementary Fig. S1*A***). Firstly, to decrease the noise content, we discarded non-informative genes based on their variability and expression levels by using the functions of *variabilityBasedfilter* and *expressionBasedfilter* of DCGL R package ^36^. Specially, for each brain area, the genes used for subsequent analysis were obtained by (*S_c_*(*exp*)∩*S_c_*(*var*)) ∪ (*S_d_*(*exp*)∩*S_d_*(*var*)), where *S_c_*(*exp*), *S_c_*(*var*) represent the gene sets filtered by the *expressionBasedfilter* and v*ariabilityBasedfilter* functions in control group, whereas *S_d_*(*exp*), *S_d_*(*var*) represent those corresponding ones in depression group, respectively. Then, the DCGs in each brain area were identified by using the I-WGCNA approach ^36, 57^. Briefly speaking, the coexpression gene networks were constructed for both the control and depression groups with the edge weight (*a_ij_*) presented by Eq. (1),

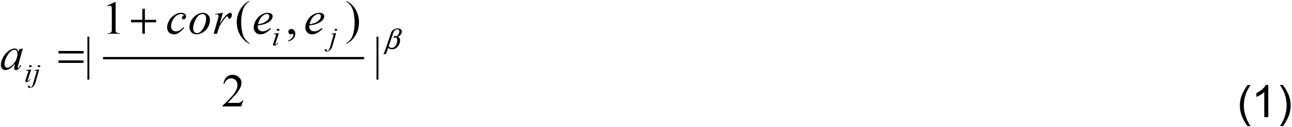

where *cor*(*e_i_, e_j_*) is the expression Spearman’s correlation coefficient (SCC) between gene *i* and *j* in control or depression group. The power value of *β* is set as 2 in order to emphasize large correlations at the expense of low correlations. For a specific gene *i*, we defined a WGCNA score (WGCNA_*i*_) to evaluate its edge weight difference between the control and depression coexpression networks (Eq. (2)),

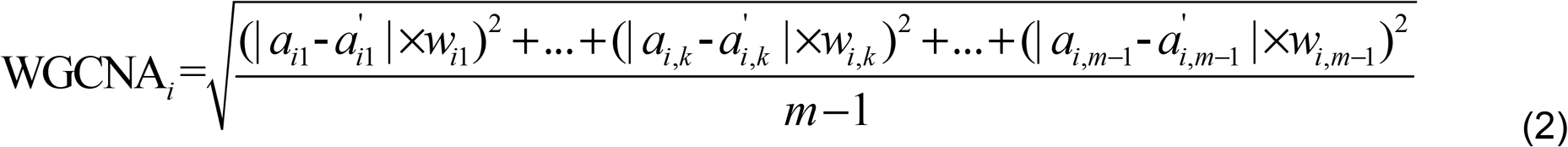

where *m* is the number of genes in a coexpression network, *a_i_* and *a*’_*i*_ are the edge weights between gene *i* with other genes in the control and depression coexpression networks, respectively. It is worth noting that we introduced a weighted factor (*w_i, k_*) in the WGCNA score instead of using the original length-normalized Euclidean distance (Eq. (3)),

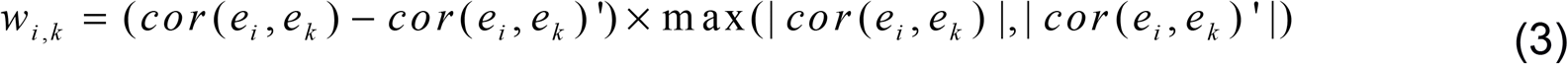

where the *cor*(*e_i_, e_k_*) and *cor*(*e_i_, e_k_*)’ are expression SCCs of gene *i* and *k* in control and depression groups, respectively. *w_i, k_* is responsible for distinguishing the importance when two pairs of genes at two groups have the same SCC diLerence. Finally, the sample permutation is repeated 10, 000 times, with P-value for each gene be estimated from an empirical null distribution. And the Benjamini-Hochberg (BH) method was used to control the false discovery rate. The pathway enrichment analysis of DCGs was performed against KEGG database using the *enrichKEGG* function in *clusterProfiler* R Package ^58^. A rank-based enrichment analysis was then carried out to detect the potential genes that regulate these enriched pathways. Specifically, for a candidate gene, we generated a gene list of its coexpression genes which are ranked according to their correlation differences calculated with Eq. (3). The relationship between the candidate gene and the enriched pathway was then determined by the enrichment score (ES) calculated by the GSEA algorithm ^59^. The randomization step was repeated 10,000 times, with the empirical *P*-value estimated by calculation of the fraction of permutations in which the random count was higher than or equal to the observed values. In addition, to control the false discovery rate, Benjamini-Hochberg (BH) method was employed.

### Statistical analyses

All data given in text and figures indicate mean ± s.e.m. For analysis of experiments with two groups, we used the parametric unpaired two-tailed Student’s t test. For analysis of experiments with three or more groups, the parametric one-way ANOVA with Dunnett’s multiple comparisons test or two-way ANOVA with Sidak’s multiple comparisons test were used. Differences were considered significant when P was < 0.05. NS = not significant, ^*^ P < 0.05, ^**^ P < 0.01, ^***^ P < 0.001.

## Supplemental Information

Supplemental Information can be found in the online version of this article.

## Acknowledgements

We thank Z. Jun for technical assistances and reagents. The research is financially supported by the National Natural Science Foundation of China (NO. U1603285 and NO. 81803960).

## Author Contributions

Z.Y.W., Y.X.F., C.H., W.X. and Y.H.W. conceived the study. Z.Y.W. assisted by Y.X.F., C.H. and C.L.Z. designed and performed majority of the experiments. C.H. and C.L.Z. generated and analyzed the data from GEO. Z.H.G. contributed to data analysis of RNA-seq. Y.X.F. and Z.H.G. oversaw all bioinformatics analyses. J.L.Z. and L.Y.C. contributed to *in vivo* surgeries and behavioral experiments. X.G.L., L.Y.C., W.W.Z., Y.Y.C., J.M.W., Y.Y. and M.J. helped with the Q-RTPCR, western blot and Immunohistochemistry experiments. S.G.S. and Z.H.Y. were responsible for the design and synthesis of lutD, Z.H.Y. also performed fluorescence spectroscopy analysis. X.T.C. screened out candidate compounds of GLO1 inhibitor. J.H.W., X.T.C. and Y.F.Y. performed molecular docking and Molecular dynamics simulation. J.L.L. and A.P.L. provided guidance and advice. Z.Z.W. provided technical assistance. Y.L. provided advice and oversaw a portion of the work. Y.X.F., Y.F.Y., Z.Y.W., C.H., and C.L.Z. wrote the manuscript. All authors discussed the results and commented on the manuscript.

## References

1 Bus, B. A. et al. Chronic depression is associated with a pronounced decrease in serum brain-derived neurotrophic factor over time. Mol Psychiatry 20, 602–608, doi:10.1038/mp.2014.83 (2015).

2 Taliaz, D. et al. Resilience to chronic stress is mediated by hippocampal brain-derived neurotrophic factor. J Neurosci 31, 4475–4483, doi:10.1523/JNEUROSCI.5725-10.2011 (2011).

3 Feder, A., Nestler, E. J. & Charney, D. S. Psychobiology and molecular genetics of resilience. Nat Rev Neurosci 10, 446–457, doi:10.1038/nrn2649 (2009).

4 Vialou, V. et al. ΔFosB in brain reward circuits mediates resilience to stress and antidepressant responses. Nat Neurosci 13, 745–752, doi:10.1038/nn.2551 (2010).

5 Hollon, N. G., Burgeno, L. M. & Phillips, P. E. Stress effects on the neural substrates of motivated behavior. Nat Neurosci 18, 1405–1412, doi:10.1038/nn.4114 (2015).

6 Shin, S. et al. mGluR5 in the nucleus accumbens is critical for promoting resilience to chronic stress. Nat Neurosci 18, 1017–1024, doi:10.1038/nn.4028 (2015).

7 Allaman, I., Belanger, M. & Magistretti, P. J. Methylglyoxal, the dark side of glycolysis. Front Neurosci 9, 23, doi:10.3389/fnins.2015.00023 (2015).

8 Shamsi, F. A., Partal, A., Sady, C., Glomb, M. A. & Nagaraj, R. H. Immunological evidence for methylglyoxal-derived modifications in vivo. Determination of antigenic epitopes. J Biol Chem 273, 6928–6936 (1998).

9 Yao, D. et al. High glucose increases angiopoietin-2 transcription in microvascular endothelial cells through methylglyoxal modification of mSin3A. J Biol Chem 282, 31038–31045, doi:10.1074/jbc.M704703200 (2007).

10 Di Loreto, S. et al. Methylglyoxal induces oxidative stress-dependent cell injury and up-regulation of interleukin-1beta and nerve growth factor in cultured hippocampal neuronal cells. Brain Res 1006, 157–167, doi:10.1016/j.brainres.2004.01.066 (2004).

11 Bierhaus, A. et al. Methylglyoxal modification of Nav1.8 facilitates nociceptive neuron firing and causes hyperalgesia in diabetic neuropathy. Nat Med 18, 926–933, doi:10.1038/nm.2750 (2012).

12 Ramasamy, R., Yan, S. F. & Schmidt, A. M. Methylglyoxal comes of AGE. Cell 124, 258–260, doi:10.1016/j.cell.2006.01.002 (2006).

13 Angeloni, C. et al. Neuroprotective effect of sulforaphane against methylglyoxal cytotoxicity. Chem Res Toxicol 28, 1234–1245, doi:10.1021/acs.chemrestox.5b00067 (2015).

14 Belanger, M. et al. Role of the glyoxalase system in astrocyte-mediated neuroprotection. J Neurosci 31, 18338–18352, doi:10.1523/JNEUROSCI.1249-11.2011 (2011).

15 Distler, M. G. et al. Glyoxalase 1 increases anxiety by reducing GABAA receptor agonist methylglyoxal. J Clin Invest 122, 2306–2315, doi:10.1172/JCI61319 (2012).

16 McMurray, K. M. J. et al. Identification of a novel, fast-acting GABAergic antidepressant. Mol Psychiatry 23, 384–391, doi:10.1038/mp.2017.14 (2018).

17 Willner, P. The chronic mild stress (CMS) model of depression: History, evaluation and usage. Neurobiol Stress 6, 78–93, doi:10.1016/j.ynstr.2016.08.002 (2017).

18 McEwen, B. S. et al. Mechanisms of stress in the brain. Nat Neurosci 18, 1353–1363, doi:10.1038/nn.4086 (2015).

19 Russo, S. J., Murrough, J. W., Han, M. H., Charney, D. S. & Nestler, E. J. Neurobiology of resilience. Nat Neurosci 15, 1475–1484, doi:10.1038/nn.3234 (2012).

20 Golden, S. A., Covington, H. E., 3rd, Berton, O. & Russo, S. J. A standardized protocol for repeated social defeat stress in mice. Nat Protoc 6, 1183–1191, doi:10.1038/nprot.2011.361 (2011).

21 Cao, X. et al. Astrocyte-derived ATP modulates depressive-like behaviors. Nat Med 19, 773–777, doi:10.1038/nm.3162 (2013).

22 Duman, R. S., Aghajanian, G. K., Sanacora, G. & Krystal, J. H. Synaptic plasticity and depression: new insights from stress and rapid-acting antidepressants. Nat Med 22, 238–249, doi:10.1038/nm.4050 (2016).

23 Lin, J. Y., Jiang, M. Y., Kan, Z. M. & Chu, Y. Influence of 5-HTR2A genetic polymorphisms on the efficacy of antidepressants in the treatment of major depressive disorder: a meta-analysis. J Affect Disord 168, 430–438, doi:10.1016/j.jad.2014.06.012 (2014).

24 Andero, R., Choi, D. C. & Ressler, K. J. BDNF-TrkB receptor regulation of distributed adult neural plasticity, memory formation, and psychiatric disorders. Prog Mol Biol Transl Sci 122, 169–192, doi:10.1016/B978-0-12-420170-5.00006-4 (2014).

25 Li, Y. et al. TrkB regulates hippocampal neurogenesis and governs sensitivity to antidepressive treatment. Neuron 59, 399–412, doi:10.1016/j.neuron.2008.06.023 (2008).

26 Atwal, J. K., Massie, B., Miller, F. D. & Kaplan, D. R. The TrkB-Shc site signals neuronal survival and local axon growth via MEK and P13-kinase. Neuron 27, 265–277 (2000).

27 Kang, H. & Schuman, E. M. Long-lasting neurotrophin-induced enhancement of synaptic transmission in the adult hippocampus. Science 267, 1658–1662 (1995).

28 Hoshaw, B. A., Malberg, J. E. & Lucki, I. Central administration of IGF-I and BDNF leads to long-lasting antidepressant-like effects. Brain Res 1037, 204–208, doi:10.1016/j.brainres.2005.01.007 (2005).

29 Tapley, P., Lamballe, F. & Barbacid, M. K252a is a selective inhibitor of the tyrosine protein kinase activity of the trk family of oncogenes and neurotrophin receptors. Oncogene 7, 371–381 (1992).

30 Jang, S. W. et al. A selective TrkB agonist with potent neurotrophic activities by 7,8-dihydroxyflavone. Proc Natl Acad Sci U S A 107, 2687–2692, doi:10.1073/pnas.0913572107 (2010).

31 Porcher, C. et al. Positive feedback regulation between γ-aminobutyric acid type A (GABA(A)) receptor signaling and brain-derived neurotrophic factor (BDNF) release in developing neurons. J Biol Chem 286, 21667–21677, doi:10.1074/jbc.M110.201582 (2011).

32 Jovanovic, J. N., Thomas, P., Kittler, J. T., Smart, T. G. & Moss, S. J. Brain-derived neurotrophic factor modulates fast synaptic inhibition by regulating GABA(A) receptor phosphorylation, activity, and cell-surface stability. J Neurosci 24, 522–530, doi:10.1523/JNEUROSCI.3606-03.2004 (2004).

33 Lo, T. W., Westwood, M. E., McLellan, A. C., Selwood, T. & Thornalley, P. J. Binding and modification of proteins by methylglyoxal under physiological conditions. A kinetic and mechanistic study with N alpha-acetylarginine, N alpha-acetylcysteine, and N alpha-acetyllysine, and bovine serum albumin. J Biol Chem 269, 32299–32305 (1994).

34 Haniu, M. et al. Interactions between Brain-derived Neurotrophic Factor and the TRKB Receptor IDENTIFICATION OF TWO LIGAND BINDING DOMAINS IN SOLUBLE TRKB BY AFFINITY SEPARATION AND CHEMICAL CROSS-LINKING. 272, 25296–25303 (1997).

35 Li, J. Z. et al. Circadian patterns of gene expression in the human brain and disruption in major depressive disorder. Proc Natl Acad Sci U S A 110, 9950–9955, doi:10.1073/pnas.1305814110 (2013).

36 Liu, B. H. et al. DCGL: an R package for identifying differentially coexpressed genes and links from gene expression microarray data. Bioinformatics 26, 2637–2638, doi:10.1093/bioinformatics/btq471 (2010).

37 Zheng, C. et al. Large-scale Direct Targeting for Drug Repositioning and Discovery. Sci Rep 5, 11970, doi:10.1038/srep11970 (2015).

38 Ru, J. et al. TCMSP: a database of systems pharmacology for drug discovery from herbal medicines. J Cheminform 6, 13, doi:10.1186/1758-2946-6-13 (2014).

39 Jafari, R. et al. The cellular thermal shift assay for evaluating drug target interactions in cells. Nat Protoc 9, 2100–2122, doi:10.1038/nprot.2014.138 (2014).

40 Kato, T. et al. BDNF release and signaling are required for the antidepressant actions of GLYX-13. Mol Psychiatry, doi:10.1038/mp.2017.220 (2017).

41 Ma, Z. et al. TrkB dependent adult hippocampal progenitor differentiation mediates sustained ketamine antidepressant response. Nat Commun 8, 1668, doi:10.1038/s41467-017-01709-8 (2017).

42 Colle, R. et al. BDNF/TRKB/P75NTR polymorphisms and their consequences on antidepressant efficacy in depressed patients. Pharmacogenomics 16, 997–1013, doi:10.2217/pgs.15.56 (2015).

43 Fava, M. Weight gain and antidepressants. J Clin Psychiatry 61 Suppl 11, 37–41 (2000).

44 Kent, J. M. SNaRIs, NaSSAs, and NaRIs: new agents for the treatment of depression. Lancet 355, 911–918, doi:10.1016/S0140-6736(99)11381-3 (2000).

45 Gu, B. et al. A Peptide Uncoupling BDNF Receptor TrkB from Phospholipase Cgamma1 Prevents Epilepsy Induced by Status Epilepticus. Neuron 88, 484–491, doi:10.1016/j.neuron.2015.09.032 (2015).

46 Petzinger, G. M. et al. Exercise-enhanced neuroplasticity targeting motor and cognitive circuitry in Parkinson’s disease. Lancet Neurol 12, 716–726, doi:10.1016/S1474-4422(13)70123-6 (2013).

47 Ishisaka, M. et al. Luteolin shows an antidepressant-like effect via suppressing endoplasmic reticulum stress. Biol Pharm Bull 34, 1481–1486 (2011).

48 Nabavi, S. F. et al. Luteolin as an anti-inflammatory and neuroprotective agent: A brief review. Brain Res Bull 119, 1–11, doi:10.1016/j.brainresbull.2015.09.002 (2015).

49 Kanai, K. et al. Therapeutic anti-inflammatory effects of luteolin on endotoxin-induced uveitis in Lewis rats. J Vet Med Sci 78, 1381–1384, doi:10.1292/jvms.16-0196 (2016).

50 Nollet, M., Le Guisquet, A. M. & Belzung, C. Models of depression: unpredictable chronic mild stress in mice. Curr Protoc Pharmacol Chapter 5, Unit 5 65, doi:10.1002/0471141755.ph0565s61 (2013).

51 West, M. J., Slomianka, L. & Gundersen, H. J. Unbiased stereological estimation of the total number of neurons in thesubdivisions of the rat hippocampus using the optical fractionator. Anat Rec 231, 482–497, doi:10.1002/ar.1092310411 (1991).

52 Kim, D., Langmead, B. & Salzberg, S. L. HISAT: a fast spliced aligner with low memory requirements. Nat Methods 12, 357–360, doi:10.1038/nmeth.3317 (2015).

53 Pertea, M. et al. StringTie enables improved reconstruction of a transcriptome from RNA-seq reads. Nat Biotechnol 33, 290–295, doi:10.1038/nbt.3122 (2015).

54 Frazee, A. C. et al. Ballgown bridges the gap between transcriptome assembly and expression analysis. Nat Biotechnol 33, 243–246, doi:10.1038/nbt.3172 (2015).

55 Huang da, W., Sherman, B. T. & Lempicki, R. A. Bioinformatics enrichment tools: paths toward the comprehensive functional analysis of large gene lists. Nucleic Acids Res 37, 1–13, doi:10.1093/nar/gkn923 (2009).

56 Paxinos, G. & Watson, C. The rat brain in stereotaxic coordinates. 6th edn, (Academic Press/Elsevier, 2007).

57 Langfelder, P. & Horvath, S. WGCNA: an R package for weighted correlation network analysis. BMC Bioinformatics 9, 559, doi:10.1186/1471-2105-9-559 (2008).

58 Yu, G., Wang, L. G., Han, Y. & He, Q. Y. clusterProfiler: an R package for comparing biological themes among gene clusters. OMICS 16, 284–287, doi:10.1089/omi.2011.0118 (2012).

59 Subramanian, A. et al. Gene set enrichment analysis: a knowledge-based approach for interpreting genome-wide expression profiles. Proc Natl Acad Sci U S A 102, 15545–15550, doi:10.1073/pnas.0506580102 (2005).

